# Optimizing immunization protocols to elicit broadly neutralizing antibodies

**DOI:** 10.1101/2020.01.04.894857

**Authors:** Kayla G. Sprenger, Joy E. Louveau, Arup K. Chakraborty

## Abstract

Natural infections and vaccination with a pathogen typically stimulates the production of potent antibodies specific for the pathogen through a Darwinian evolutionary process known as affinity maturation. Such antibodies provide protection against reinfection by the same strain of a pathogen. A highly mutable virus, like HIV or influenza, evades recognition by these strain-specific antibodies via the emergence of new mutant strains. A vaccine that elicits antibodies that can bind to many diverse strains of the virus – known as broadly neutralizing antibodies (bnAbs) – could protect against highly mutable pathogens. Despite much work, the mechanisms by which bnAbs emerge remain uncertain. Using a computational model of affinity maturation, we studied a wide variety of vaccination strategies. Our results suggest that an effective strategy to maximize bnAb evolution is through a sequential immunization protocol, wherein each new immunization optimally increases the pressure on the immune system to target conserved antigenic sites, thus conferring breadth. We describe the mechanisms underlying why sequentially driving the immune system increasingly further from equilibrium, in an optimal fashion, is effective. The optimal protocol allows many evolving B cells to become bnAbs via diverse evolutionary paths.

**Significance Statement:** The global health burden could be substantially alleviated by the creation of universal vaccines against highly mutable pathogens like HIV and influenza. Broadly-neutralizing antibodies (bnAbs) are encouraging targets for such vaccines, because they can bind to diverse strains of highly mutable pathogens. BnAbs typically develop only rarely upon natural infection, after the immune system has been exposed to many mutated versions of a pathogen. Thus, sequentially administering multiple different pathogen-like proteins (antigens) is a promising strategy to elicit bnAbs through vaccination. However, it remains unclear how best to design and administer these antigens. We explore this matter using physics-based simulations, and provide new mechanistic insights into antibody evolution that could guide the creation of universal vaccines against highly mutable pathogens.

## Introduction

Successful vaccines stimulate immune responses that can protect the host from infection by a specific pathogen. Such a vaccine is usually unable to protect against the diverse circulating strains of highly mutable pathogens. Examples include viruses like HIV and HCV^1^. Such a vaccine can also not serve as a universal vaccine against variant strains of the influenza virus that arise annually. Creating an effective universal vaccine against highly mutable pathogens will likely require a novel approach to vaccine design.

Successful prophylactic vaccines induce the immune system to generate antibodies (Abs) that can bind to and neutralize the pathogen. The process that governs Ab production upon pathogen (or antigen, Ag) encounter/vaccine stimulation is a stochastic Darwinian evolutionary process called affinity maturation (AM)^2,3^. First, a few B cells become activated upon binding of their B cell receptors (BCRs) to the Ag with sufficient affinity. Activated B cells can then seed microstructures called germinal centers (GCs) in lymph nodes. During AM, B cells proliferate, and upon induction of the activation-induced cytidine deaminase (AID) gene, mutations are introduced into the BCR at a high rate via a process known as somatic hypermutation. The B cells with mutated receptors then interact with the Ag displayed on the surface of GC-resident follicular dendritic cells (FDCs) and attempt to bind and internalize the Ag. B cells with BCRs that bind to the Ag with higher affinity are more likely to internalize Ag. The internalized Ag is then processed and displayed on the surface of B cells as peptide-major histocompatibility complex (MHC) molecules. B cells that display peptide-MHC molecules then compete with each other to interact with T helper cells, productive binding of whose T cell receptors (TCRs) to peptide-MHC molecules delivers a survival signal. B cells that bind more strongly to the Ag on FDCs are likely to internalize more Ag and thus likely to display more peptide-MHC molecules on their surface, and are therefore more likely to be positively selected^4^. B cells that do not bind the Ag strongly enough or do not receive T cell help undergo apoptosis^5,6^. A few positively selected B cells exit the GC and differentiate into Ab-producing plasma cells and memory B cells, while the majority are recycled for further rounds of mutation and selection^3,7^. Upon immunization with a single Ag, as cycles of diversification and selection ensue, Abs with increasingly higher affinity for the Ag are thus produced^8^.

One promising strategy for the development of an effective vaccine against HIV is to induce the immune system to generate broadly-neutralizing antibodies (bnAbs) that can neutralize diverse strains of mutable pathogens. BnAbs that can neutralize most HIV strains *in vitro*^9–12^ and diverse influenza strains have been isolated^13,14^. These Abs target regions or epitopes on the surface of pathogenic proteins that contain amino acid residues that are relatively conserved because they are key to the virus’ ability to propagate infection in human cells. One possible strategy to generate bnAbs is to vaccinate with an immunogen (an immune response-eliciting Ag) containing only these conserved residues. This tempting solution is impractical, however, because the Abs thus generated cannot learn how to bind to the conserved residues in the molecular context in which they are presented on the pathogen surface. For example, an important target of bnAbs against HIV is the CD4 binding site on the virus’ spike, and the conserved residues therein are surrounded by glycans and variable residues that, due to the three-dimensional nature of the HIV spike proteins, partially shield B cell receptors (BCRs) from binding to the conserved residues. In fact, mutations in variable antigenic residues can insert loops that further hinder access to conserved residues^15^. Similarly, for influenza, the receptor binding site on the head of its spike is surrounded by variable residues^13^, and another conserved region in the stem of the spike is sterically difficult to access because of the high density of spikes on the influenza virus’ surface^16^. A potentially promising vaccination strategy that may elicit bnAbs is to first administer an Ag that can stimulate germline B cells that can target the pertinent conserved residues, followed by immunization with variant Ags that share the same amino acids at conserved positions, but diverse amino acids at the surrounding variable positions presented in the same molecular context as in the real pathogen’s spike. A deep understanding of how Abs evolve in such a vaccination setting is required to be able to guide the design of optimal vaccination strategies that can efficiently stimulate the production of bnAbs^17–21^ against different highly mutable pathogens in diverse individuals. Developing this knowledge also presents an interesting challenge at the intersection of immunology, evolutionary biology, and biophysics.

In cases where bnAbs develop in HIV-infected individuals, their emergence typically occurs only after several years of infection, during which the immune system of the infected individual has been exposed to many different antigenic strains of the rapidly-mutating virus^22–24^. BnAbs against influenza have similarly been observed to emerge in rare individuals upon exposure to antigenically distinct strains of the virus^13,14^. These data suggest that being selected by different, but somewhat related, Ags promotes the evolution of bnAbs. The diverse infecting Ags serve as selection forces that shape Ab evolution during AM. Can properly designed vaccination protocols using variant Ags that share conserved residues but differ in the variable regions result in AM that elicits bnAbs efficiently in diverse individuals? If so, a strategy for developing vaccines against highly mutable pathogens would be available. This tantalizing possibility has led to a great deal of research directed toward achieving this goal.

A number of advances have been made in this regard, and we note just a few that have resulted from work focused on eliciting bnAbs against HIV. Strategies and immunogens have been designed to activate the correct germline B cells that can target conserved residues on the HIV viral spike, which have the potential for developing into bnAbs upon subsequent immunization with variant Ags^25–27^. Computational studies have described how the variant Ags can serve as conflicting selection forces because a BCR/Ab that binds well to one of the variant Ags is unlikely to bind well to another variant Ag unless relatively strong interactions with the shared conserved residues evolve. The presence of such conflicting selection forces has been termed “frustration”, too much of which has been shown to result in substantial B cell death/GC extinction^28,29^. Studies have suggested that, in some instances, sequential immunization with variant Ags may elicit bnAbs more efficiently than a cocktail of the same Ags^28,29,26,29–34^. These studies also highlight the importance of separating the conflicting selection forces (or sources of frustration) over time to minimize significant B cell death, and of using multiple immunizations to help the developing Abs acquire mutations that focus their binding on to the conserved residues, and thus confer breadth. It has also been suggested that cocktails can potentially be optimally designed to promote bnAb formation while minimizing cell death by manipulating the vaccine dose and mutational distance between Ags^28^, thus identifying these variables as two further sources of frustration that can influence the outcome of AM.

Despite this progress, several questions remain to be answered to devise immunogens and vaccination protocols that can efficiently induce bnAbs. In particular, 1) which Ags should be used as immunogens (number of Ags and their sequences/compositions), and 2) how should they be administered in time? Given the high diversity of the variable residues as well as the broad range of possible BCR-Ag binding sites, the number of Ags that share conserved residues and the number of different immunization protocols are too great to test all possible combinations of these Ags and protocols with experiments in animal models. It is, however, possible for computational/theoretical studies to elucidate the effect of these variables and evaluate the outcomes in terms of both the quality (mean breadth) and quantity (titers) of the produced Abs. These models could provide new insights into how immunogens and vaccination protocols influence bnAb evolution and thus help to guide the design of an Ab-based vaccine against highly mutable pathogens. Such studies can also shed light on fundamental questions in immunology and biophysics.

In this study, we used a computational model to explore how different sets of variant Ags administered sequentially at different concentrations influences the evolution of bnAbs by AM. By quantifying frustration and characterizing its impact on AM, we predict that Ags and vaccination protocols that result in a temporally increasing level of frustration on GC reactions promotes optimal bnAb responses. This can be realized, for example, by decreasing the Ag concentration for each new immunization, or by immunizing with increasingly dissimilar Ags. Our results further indicate that an intermediate amount of frustration during each new immunization is optimal. The optimum is defined by the highest level of frustration allowed that does not result in extensive GC collapse/cell death. We describe the mechanisms underlying these results, which highlight that an appropriate level of diversity among B cells needs to evolve early on. This optimal level of diversity allows B cells to subsequently acquire mutations that confer breadth by diverse evolutionary trajectories.

## Results

### Description of the computational model of affinity maturation

#### Overview

We built a simple computational AM model of B cells in the presence of different Ags that can be introduced at several time points. We simulate the processes that occur during AM using a stochastic model, as is appropriate for an evolutionary process. The set of rules that define AM are derived from experimental studies of AM with a single Ag^3,35–37^, and these serve as instructions that are executed by the computer. The goal of this model is not the quantitative reproduction of existing experiments, but to provide mechanistic insights into how the nature of variant Ags and immunization protocols influence the development of Ab breadth and bnAb titers. While our model represents the steps of AM, it does not explicitly account for B cell migration within a GC or employ an atomistically detailed representation to compute the free energy of binding of a BCR to Ags. The model is based on those of previous works^28,29,38^, but there are a number of new features that are described below. All parameters that will be discussed are listed in Table S1.

#### Binding Free Energy

As in the models described by (Wang et al. 2015, Luo et al. 2015), the BCR paratope and the Ag epitope of interest, which we will from here on refer to as BCR and Ag, respectively, are both represented by a string of residues^29,39^. The BCR and the Ag have identical lengths (46 residues total; 28 variable and 18 conserved residues), so the BCR residue at position *k* binds to the antigenic residue located at the same position *k*. For the BCR, the identity of each residue is designated by a number, whose value was sampled from a continuous and bounded uniform distribution; variable residue values were bounded between −0.18 and 0.90 (bounds change after AID gene turns on (see Table S1)), and conserved residue values were bounded between 0.3 and 0.6 (see following section for more details). Similarly, each Ag residue is designated by a number. The value of this number corresponding to a conserved Ag residue is always +1, while the value of a variable residue can be either +1 (non-mutated) or −1 (mutated). The binding free energy depends on the strength of the interactions between the BCR and the Ag. We model this binding free energy as a sum of all pairwise interactions between the BCR and the Ag, estimated as the product of the values that describe the identity of each BCR residue and its analogous antigenic residue. Thus, the binding free energy is:

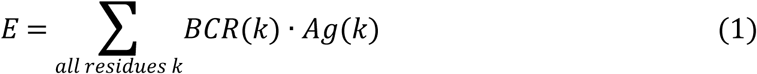

More positive binding free energies correspond to stronger affinities. This simple expression for the binding free energy is further embellished to account for the effects of three-dimensional conformations of the BCR and Ag as described in context below. The absolute value of the binding free energy is arbitrary as it is determined only up to an additive constant. All binding free energies in our model are expressed in units of the thermal energy (*k*_*B*_*T*). We assume that a binding free energy of 9 *k*_*B*_*T* is the minimum requirement for Ag binding and therefore only B cells that bind to an Ag with a binding free energy above this threshold can seed a GC. The value of this threshold binding free energy sets the scale for our calculations, and we do not expect qualitative results to depend upon this particular choice.

#### Germline-targeting and activation of a GC

The number of potential BCR/Ab binding sites on the HIV envelope that do not contain conserved residues far exceeds those that could be bnAb epitopes^40^. In the repertoire of naïve B cells bnAb precursors are rare^27^, and thus the precursor frequency of germline B cells that target epitopes that do not contain conserved residues is much higher. Thus, it is unlikely that bnAb precursors will get activated during a natural infection^41^. Similar considerations apply to influenza bnAbs, where there are many more highly variable hemagglutinin (HA) head epitopes than those that can potentially be epitopes for bnAbs. Immunization experiments for HIV try to remedy this challenge by priming with an Ag that focuses immune responses on conserved residues (Fig. 1). That is, immunizing with Ags designed to target specific germline (GL) BCRs is a key first step to eliciting bnAbs^25,27,33,42^. Subsequently, variant Ags that mimic the viral spike are used as immunogens (Fig. 1). Our model mimics the priming to activate the appropriate GL BCRs by seeding the first GCs, upon immunization with variant Ags, with a pool of B cells that we assume has been generated by a prior, GL-activating immunization, and thus bind to epitopes containing conserved residues (e.g., the CD4 binding site for HIV). Mimicking the BCRs produced through GL-targeting experiments^25^, we chose the residues of the BCRs that bind to the conserved residues of the Ag to be slightly biased towards positive values, reflecting that they favor these residues because they were activated initially by the GL-targeting immunogen. Since no specific selection force was imposed on the residues of these BCRs that bind to the variable residues of the Ags, these were chosen to be highly variable among the seeding B cells. Experiments have shown that the number of seeding B cells varies between a few to a hundred cells^37^. We chose to seed GCs with 10 cells; changing this number did not affect our qualitative results. We assume that the GL-targeting immunogen residues are all +1s. This is just a reference sequence, and a different choice would lead to a linear shift in the mutational distances that we calculate for the variant Ags used for subsequent immunizations, and thus would have no effect on the qualitative results we report.

**Fig. 1.**
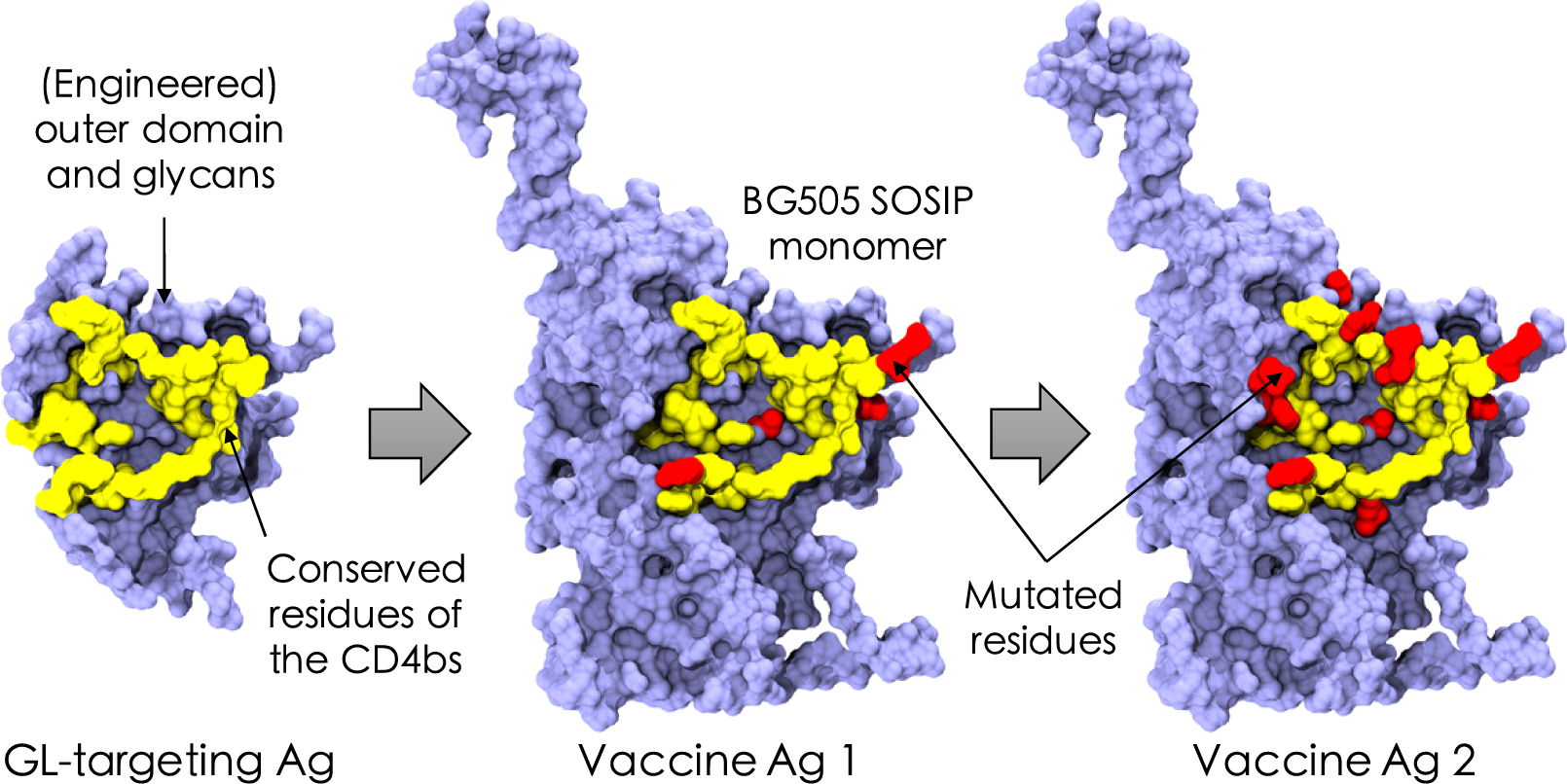
Schematic of the *in silico* immunization scheme. The germline (GL)-targeting antigen (Ag) was inspired by the eOD-GT8 construct designed to target precursor naïve B cells against the CD4 binding site (CD4bs) of HIV^27^. Conserved residues of the CD4bs are schematically depicted in yellow, example mutated variable residues are shown in red, and the surrounding residues are shown in blue. Visual Molecular Dynamics (VMD)^52^ was used to construct the images from PDB code 5fyj^53^.

#### B cell expansion and somatic hypermutation in the dark zone

The B cells expand in the dark zone of the GC without mutation or selection for a week, reaching a population size of 5,120 cells. After the initial expansion of the B cell population, the AID gene turns on and mutations are introduced into the BCRs with a probability determined by experiments: each B cell of the dark zone divides twice per GC cycle (4 divisions per day)^43^, and mutations appear at a high rate (0.14 per sequence per division)^36^. These mutations are known as somatic hypermutations (SHMs) and can have various effects on the fate of a B cell^44,45^. Recent evidence suggests that B cells that internalize more Ag divide more times^4^, but this effect is not included in our simulations. Experiments have shown that SHMs are lethal 50% of the time (for example, by making the BCR unable to fold properly), are silent 30% of the time (due to redundancy of the genetic code; i.e., synonymous mutations), and modify the binding free energy 20% of the time^36^. The energy-affecting mutations are more likely to be detrimental; experiments have determined that across all protein-protein interactions, only ~5 to 10% of energy-affecting mutations strengthen the binding free energy^46^.

For mutations that affect the binding free energy, a particular residue on the BCR is randomly picked to undergo mutation. The change in the strength of binding between this BCR residue and the complementary antigenic residue is sampled from a bounded lognormal distribution whose parameters are chosen to approximate the empirical distribution of changes in binding free energies of protein-protein interfaces upon single-residue mutation^43^. Thus, the random change in binding free energy Δ*E* is drawn from the following probability distribution:

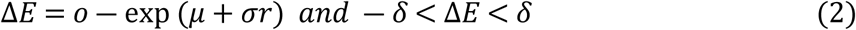

where *r* is a standard normal variable with mean zero and standard deviation equal to one. Here *o* is a shift parameter, which is needed to center the lognormal distribution properly with respect to zero, and μ and σ are the mean and standard deviation of the lognormal distribution, respectively. The parameters of the lognormal distribution (*o*, μ, σ and *r*) are set so that only 5% of mutations increase the binding free energy and so that the tail of harmful mutations fits the distribution obtained by experiments. The additional parameter δ limits the effect of a single-residue mutation, chosen to prevent the B cell population from succeeding too fast and exceeding the average time for the GC population to reach its initial size after vaccination or infection with a single Ag (see *Parameters* section).

#### Steric and conformational effects on BCR-Ag interactions

The effects of Ag residues that shield the conserved residues from being accessed easily by the BCR are incorporated into the model to account for the three-dimensional nature of Ag-BCR interactions. Mutations in variable residues can insert loops that hinder access to conserved residues^15^: greater binding to loop residues reduces access to conserved residues, and vice versa. The insertion or deletion of a loop can drastically change the binding free energy^47^. In particular, a new interaction with a loop residue can completely prevent BCR binding to non-loop residues and greatly lower the binding free energy. We mimic this effect as follows. If mutation of a BCR residue results in a stronger interaction with a variable antigenic residue, the interaction between a randomly chosen BCR residue that binds to a conserved antigenic residue is proportionally decreased by a factor α (Eqn. 3). Similarly, a weaker interaction with a variable residue leads to a proportional increase in the binding to a conserved residue. The boundary for the free energy change due to a loop *δ*_*L*_ is chosen to be the same as that for a single-residue mutation (*δ*). This feature in our model favors the emergence of mutations that alleviate steric effects because it strengthens binding to conserved residues, and this aids binding to multiple variant Ags. Mathematically, this effect is represented as follows:

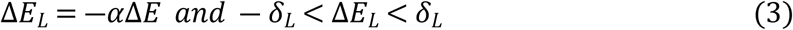

See SI for an explanation of the choice of the parameter, α.

#### Selection in the light zone

After SHM, the mutated B cells migrate to the light zone of the GC, where selection takes place through competition for binding to Ag and receiving T cell help^4^. B cells with the greatest binding free energy for the Ag presented on FDCs have a better chance to bind that Ag and present its peptides on their MHC molecules to receive T cell help. Productive interactions with T helper cells results in positive selection, and otherwise B cells undergo apoptosis^48–50^. We model this biology with a two-step selection process. First, each B cell successfully internalizes the Ag it encounters with a probability that grows with the binding free energy and then saturates, following a Langmuir form (Eqn. 4). Only the B cells that successfully internalize Ag can then go on to the second step. The surviving B cells are ranked according to their binding free energy, a proxy for the concentration of peptide-MHC molecules that they display, and only the top 70% are selected. This selection probability, as well as the parameter *e*_*scale*_ in Eq. 4 below (pseudo inverse temperature, chosen to be 0.08 k_B_T^−1^), were manually adjusted to fit with experimental observations of GC dynamics upon single-Ag administration (see Parameters section for more details)^37^. The probability of B cell *j* internalizing Ag *i* depends on the binding free energy 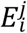 and the concentration *c*_*i*_ of that Ag, as well as the energy scale *e*_*scale*_ and the activation threshold *E*_*act*_ (Eqn. 4):

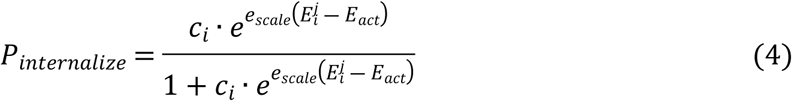

#### Recycling to the dark zone, exit for differentiation, and termination of the GC reaction

Most B cells that are positively selected are recycled for further rounds of mutation and selection, while a few randomly selected B cells exit the GC to mimic differentiation into memory and Ab-producing plasma cells^7^. As a proxy for the fact that all the Ag will be consumed by internalization if a sufficient number of B cells successfully mature, the GC reaction is terminated if the number of B cells exceeds the initial GC population size of 5,120 cells. In addition, to reflect Ag degradation over time, termination occurs when the number of cycles before the GC recovers its initial size exceeds 250 (125 days). The GC reaction also ends if all B cells die.

#### Seeding a new GC with previously-produced memory B cells after a new immunization

Every memory B cell that exits a GC is in circulation and could be reactivated by a second exposure to the Ag that initially triggered its development. Memory B cells can also then seed a new GC upon immunization with a second Ag that is different from the first Ag but shares conserved residues. If there are many other epitopes on the Ag that do not contain conserved residues, naïve B cells that bind to these epitopes could also seed GCs. However, we assume that memory B cells outnumber these naïve B cells, or that the second Ag is given sufficiently soon after the first immunization that it joins ongoing GCs with the first Ag almost depleted. Thus, we consider that only memory B cells undergo AM upon the second immunization. Ten memory cells are randomly chosen to seed a new GC.

#### Parameters

Our model depends on several parameters (Table S1). Many of these parameters represent specific biological quantities while the remaining parameters arise because of the coarse-grained nature of the model. The biological quantities can be split into those that have already been assessed by experiments – such as the SHM rate and the fate of mutations – and those that have yet to be measured experimentally. We estimated the latter type of biological quantities by making reasonable guesses. Experiments have so far focused on AM in the case where only one Ag is present, thus we fit the parameters so that simulation results are in qualitative agreement with experiments with a single Ag. Using this methodology, we fit the following parameters that control both the growth of the B cell population and the properties of the produced Abs: 1) *F*_*help,cutoff*_, the fraction of B cells that receive T cell help after binding Ag; 2) *e*_*scale*_, the factor multiplying k_B_T in the equation for *P*_*internalize*_ (Eq. 4); 3) *P*_*recycle*_, the probability to be recycled from the light zone to the dark zone; 4) *c*_*i*_, the concentration of the antigen(s); and 5) δ, the maximum value of a single-residue mutation. *F*_*help,cutoff*_, *e*_*scale*_, *P*_*recycle*_, *c*_*i*_, and δ are adjusted so that the population decreases sharply at the beginning of AM until a few beneficial mutations appear and allow survival of a few B cells. Then the population plateaus for about 20 days until enough good mutations accumulate to increase the binding free energy dramatically. The population then rises quickly until it reaches the initial size of 5,120 cells, approximately 60 days post immunization. The affinity of the Abs produced during AM with a single Ag accumulate about 10 mutations and see their binding affinity grow by at least 1,000-fold^37^. With our chosen parameters (Table S1), our model reproduces these features well.

#### Output/Analysis

For each GC reaction we record the total number of B cells over time, their binding affinities to a test panel of Ags to define the breadth of each Ab (see below), the number of energy-affecting mutations acquired by each BCR, the value of each residue (identity of pseudo amino acid) of every BCR and the binding free energy of the BCR and the Ag it interacted with in the light zone during the GC reaction. Due to the stochastic nature of AM, we executed many simulations under identical conditions (see below) and aggregated the results to obtain meaningful statistics. As in the AM model described by^38^, we group B cells into sets of functionally identical B cells, called B cell clones. All cells of a clone have the same properties, including binding free energies, number of mutations, and breadth. The size of a clone varies with time and its evolutionary trajectory.

In order to determine the breadth of coverage of each Ab, we compute the binding free energy of each clone against an artificial panel of 100 Ags different from those that the clone matured against. We found that increasing the size of the panel to 1,000 Ags produced little-to-no change in our breadth estimates. These panel Ags share the conserved residues but the value of the variable residues has equal chance to be +1 or −1. We compute the binding free energies of the clone for each panel Ag and the breadth is calculated as the fraction of panel Ags for which the clone binds with a binding free energy above a certain threshold *E*_*th*_ (Eqn. 5). The threshold was set to 12 k_B_T so that B cells produced by single-Ag immunization had a breadth of 0. This criterion reflects the fact that the mutations required to confer breadth are unlikely to evolve appreciable numbers in a single immunization, as implied by the fact that naturally evolving bnAbs usually emerge several years after infection, and sequential immunizations with variant Ags results in the evolution of bnAb-like Abs in mice^29,30^.

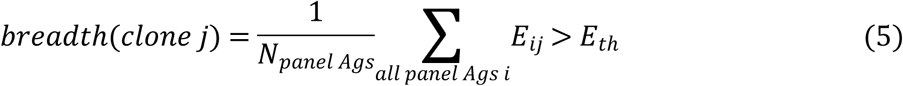

A lower threshold would define Abs generated from immunization with a single Ag as “broad” while a higher threshold would result in few or no Abs that would ever be defined as broad. In fact, the affinity of B cells for a single Ag can increase during AM by up to a few thousand-fold but above a certain value the affinity saturates. The goal of this boundary may be to safeguard against potential autoimmune responses^51^. We ran 1,000 GC trials per vaccination setting and calculated a mean breadth of all clones produced across all 1,000 GC trials (Eqn. 6). We then averaged this mean clonal breadth across multiple simulations for each vaccination protocol studied.

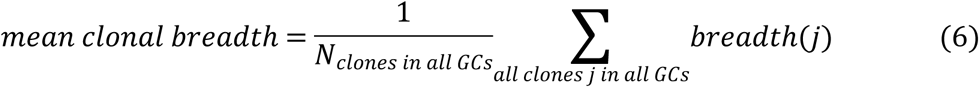

Apart from breadth, the quantity of high-breadth Abs produced (bnAb titers; clonal breadth > 0.8) by a vaccination protocol is a key metric of success. This is because low titers of bnAbs are unlikely to confer protection. A universal and efficient vaccine would likely need to elicit sufficiently high titers of high-breadth Abs. Thus, apart from the mean clonal breadth for a given vaccination setting, we also compute the average bnAb titers/GC (referred to simply as the bnAb titers/GC). We ran 1,000 GC trials per vaccination protocol and combined the outcome of all 1,000 trials in our calculations (Eqn. 7). As with the mean clonal breadth, we then averaged this value across multiple simulations for each vaccination protocol.

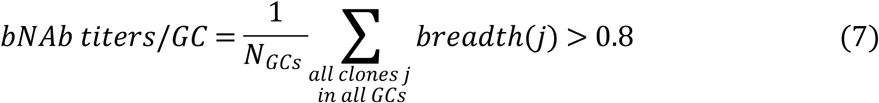

Due to the large number of GCs we analyzed (1,000) for each vaccination setting, our statistical results are robust (Fig. S1).

### An optimal level of frustration maximizes bnAb titers

We studied the situation where there are two sequential immunizations following the consequences of an assumed GL-targeting immunization (Fig. 1; see previous section on GL-targeting). Past work has shown that variant Ags subject B cells undergoing AM to conflicting selection forces, an effect referred to as frustration because these forces can, under some circumstances, frustrate the evolutionary process and lead to GC collapse. Past work has also shown that the extent of frustration can be modulated by changing the concentration of the Ags and the mutational distances between them. We first performed simulations of a single immunization with one Ag. In the results described below, we study the effects of the Ag concentration (c1) and the mutational distance (d1) between the vaccine Ag and previously-administered GL-targeting Ag (sequence of all +1s).

We find that the mean clonal breadth increases with increasing frustration (Fig. 2A-B), arising from changes in sequence (as d1 is increased) and concentration (as 1/c1 is increased). The bnAb titers/GC goes through a maximum with changes in frustration (Fig. 2C-D). The origins of these results can be understood as follows. At low frustration (Fig. 2: d1≤6 (left), c1≤1.25 (right)), the GC success rate is high (Fig. 2E-F). However, due to the ease with which the GC B cells can succeed at being positively selected, there is too little time to acquire mutations before the successful B cells consume all the Ag and the GC reaction ends (Fig. 2G-H), which results in a low diversity of clonal breadth (Fig. 2I-J). As a consequence, essentially none of the B cell clones can acquire much breadth. This is illustrated by graphs showing the variation in breadth among the resultant clones (Fig. 3A-D). Thus, the bnAb titers/GC is essentially zero.

**Fig. 2.**
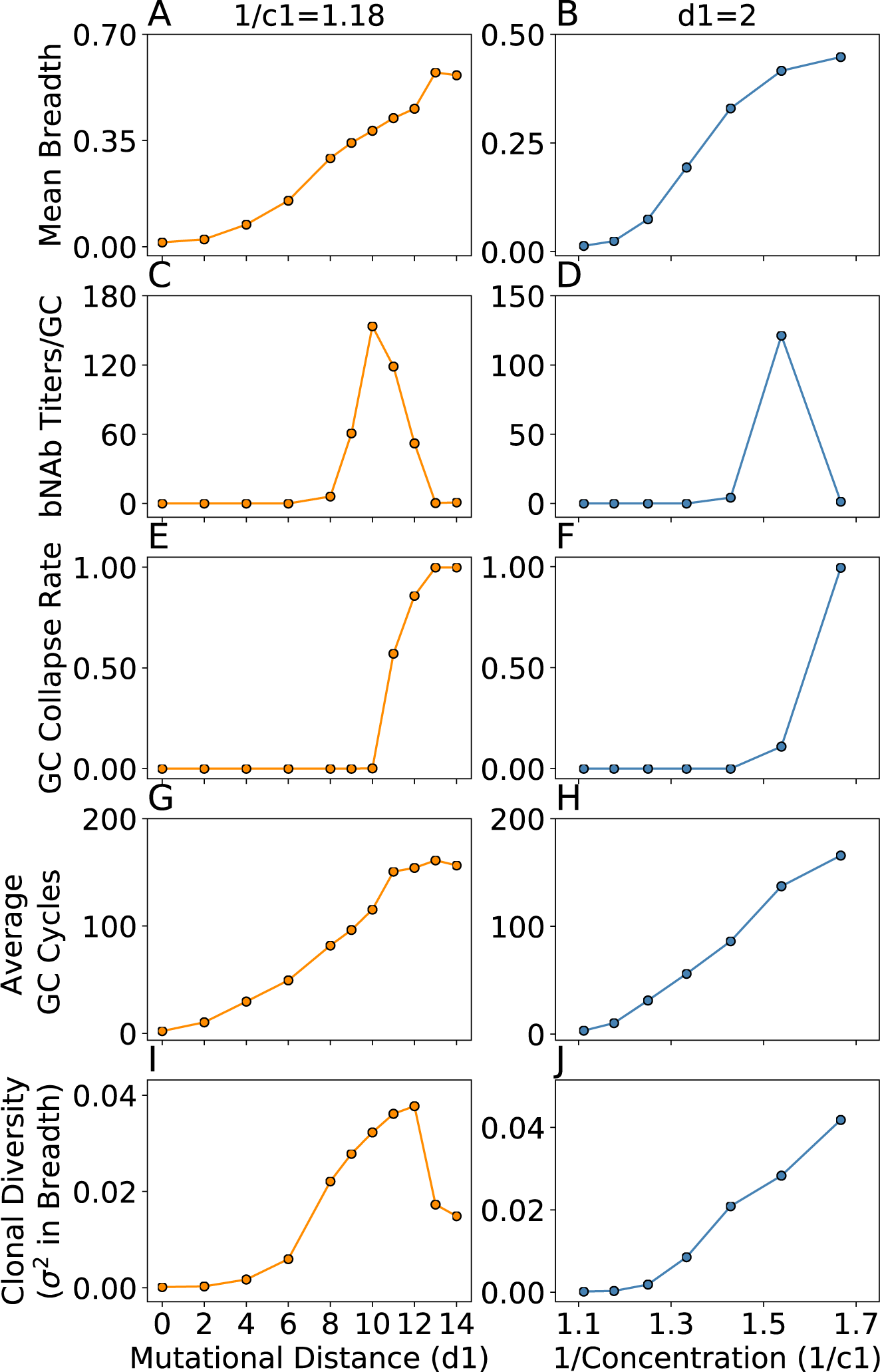
The outcomes of AM after a single vaccine immunization (second overall immunization after GL-targeting) are presented above after changing mutational distance d1 at constant concentration 1/c1 (left), and concentration 1/c1 at constant d1 (right). The measured parameters from the simulations are mean clonal breadth (A-B), bnAb titers/GC (C-D), fraction of collapsed GCs (E-F), average number of GC cycles (G-H), and clonal diversity (variance in breadth; I-J). Gray dotted lines indicate the point beyond which GCs are unlikely to be seeded due to a low Ag-BCR binding affinity. Error bars represent the standard deviation of the mean across multiple simulations (n=1,000 GCs). Note that some error bars are too small to be visible, but are included on all points and are largest where some GC collapse occurs (introducing much stochasticity into the data), or at the highest levels of frustration where few statistics could be obtained altogether.

**Fig. 3.**
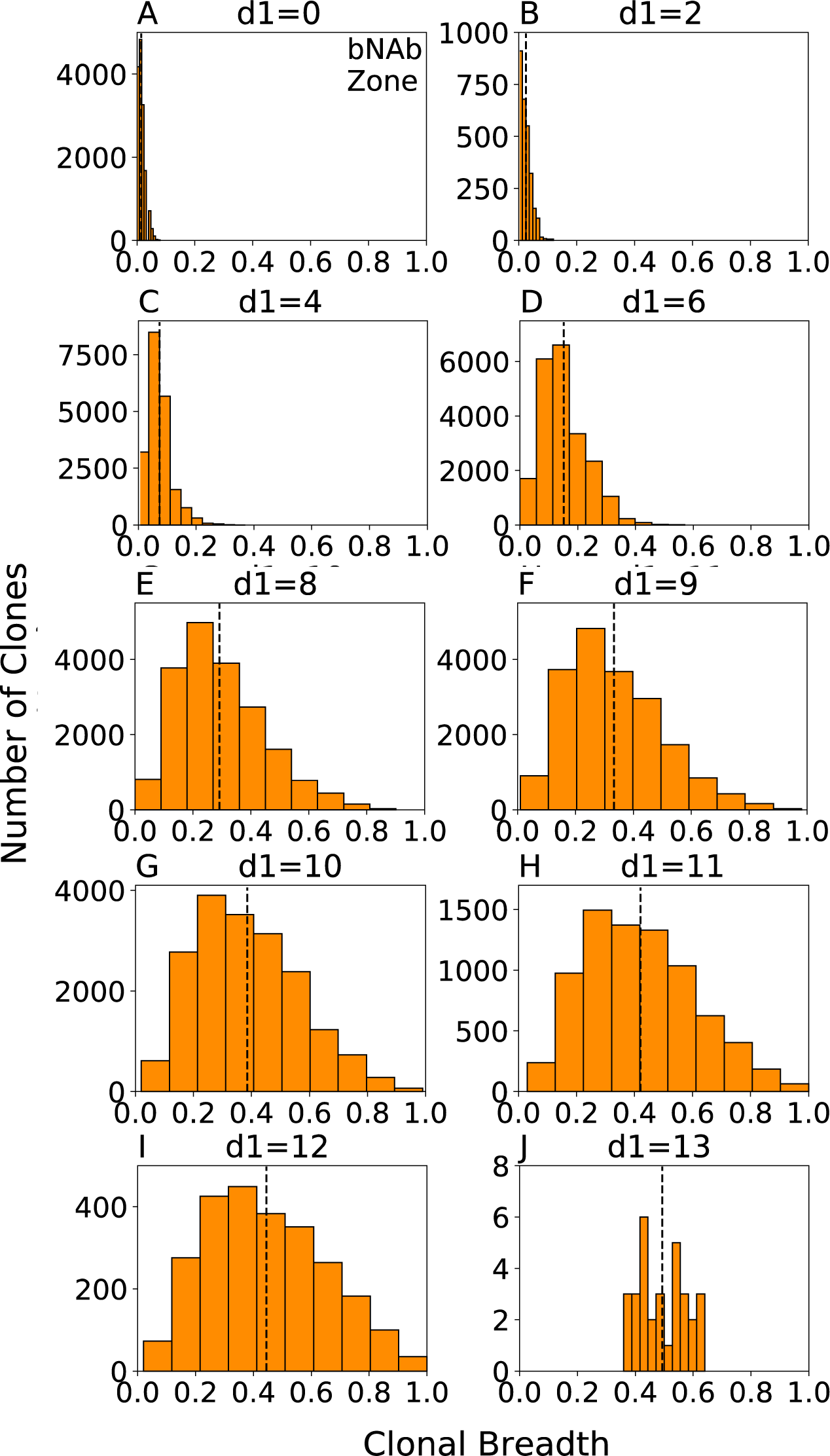
Clonal breadth distributions characterize the diversity among clones across different GCs, corresponding to the points in Figs. 2A, C, E, G, and I. A shaded blue region in each plot indicates the threshold above which BCRs are considered to have acquired breadth equivalent to a bnAb (breadth > 0.8). The amount of overlap with this region (the “bnAb zone”) is indicative of the overall bnAb titers produced at the given vaccination condition indicated above each plot. A black dashed line indicates the mean clonal breadth at that particular vaccination setting. Results are plotted for all clones produced in all 1,000 GC reactions at each vaccination setting.

As the level of frustration increases, the mean breadth and bnAb titers/GC begin to increase (Fig. 2: 6<d1≤10 (left), 1.25<c1≤1.54 (right)). This is because positive selection of B cells is less likely and so the GC reaction can continue longer without Ag depletion, enabling BCRs to acquire more mutations and increase their breadth. Thus, a more diverse population of B cells evolve with some BCRs acquiring reasonable breadth (Fig. 3E-G). As the frustration increases further, a point is eventually reached (d1=10, 1/c1=1.54 in Fig. 2C-D) beyond which the likelihood of B cells being positively selected decreases sharply, resulting in extensive GC extinction. Any additional increase in frustration results in a decline in the bnAb titers/GC. At the level of frustration corresponding to the peak in the bnAb titers/GC, the number of GC cycles is as high as possible to allow for the most time to make affinity-increasing mutations without incurring major GC extinction. In other words, more time for AM to take place – enabled by more frustration – is beneficial for increasing the mean breadth, but ultimately hurts bnAb titers because of decreased GC success rates. Thus, an intermediate level of frustration optimizes the time for AM to take place, creating the best balance between the resultant mean breadth and bnAb titers/GC. Note that as bnAb titers decline, breadth can still increase, up to a point, because diverse clones can still evolve (Fig. 3).

Ultimately, when frustration becomes very high, all that happens is GC extinction. For example, beyond a mutational distance of 12, the binding free energy between the seeding B cells and the administered Ag would likely be too weak to seed an actual GC. Yet, such high frustration settings might still be achievable in an experimental setting by manipulating concentration, and so we proceeded to simulate this regime (Fig. 2: 1/c1>1.54 (right)). The results for mutational distances beyond 12 were included purely for completeness (Fig. 2: d1>12 (left)), and were achieved by simply not setting a threshold Ag-BCR binding affinity for starting the GC reaction. At these high levels of frustration, we find that the changes in the number of cycles and mean breadth stall, almost all GCs collapse rapidly, and the very small population of B cells that survives contains no bnAbs (requires lower concentrations than explored/presented to see in Fig. 2J). Under these conditions, there are only rare evolutionary pathways to success (explored later in more detail).

### A single metric for predicting antibody properties after multiple immunizations

Our results thus far show that properties of the clonal population vary in similar ways with the level of frustration (Fig. 2), either through modulating the mutational distance between Ags (d1) or concentration (c1). Additionally, our results suggest a tradeoff in frustration exists between these two variables in the determination of mean breadth. For example, a mean breadth of 0.42 can be achieved with either (d1, 1/c1) = (11, 1.18) in Fig. 2A, or with a decreased mutational distance but increased inverse concentration of (d1, 1/c1) = (2, 1.54) in Fig. 2B. We thus hypothesized that these two sources of frustration could be combined into a single metric for predicting breadth, allowing for a simpler comparison of different temporal patterns of immunization. To test this hypothesis, we chose a simple linear combination of the two individual sources of frustration (Eqn. 8).

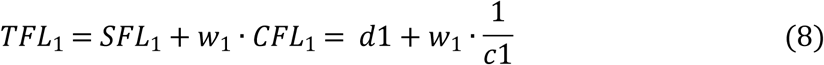

TFL refers to the total frustration level (subscript refers to the first vaccine immunization), due to the combined effects of the frustration originating from the mutational distance of the variant Ag from the GL-targeting Ag (SFL, sequence frustration level) and Ag concentration (CFL, concentration frustration level). Note that the CFL has been written as 1/c1 because administering less Ag results in a higher level of frustration during the GC reaction. The parameter, *w*, describes the relative weight of the two contributions to the total level of frustration. By fitting the value of *w*, we find that all the simulation results described in the preceding section collapse onto a single curve that is highly predictive of the resultant mean clonal breadth after immunization with a single Ag (Fig. 4A). Furthermore, with the same value of *w*, the bnAb titers/GC exhibits a maximum at an intermediate TFL_1_ (Fig. 4B), thus encapsulating the results in Fig. 2.

**Fig. 4.**
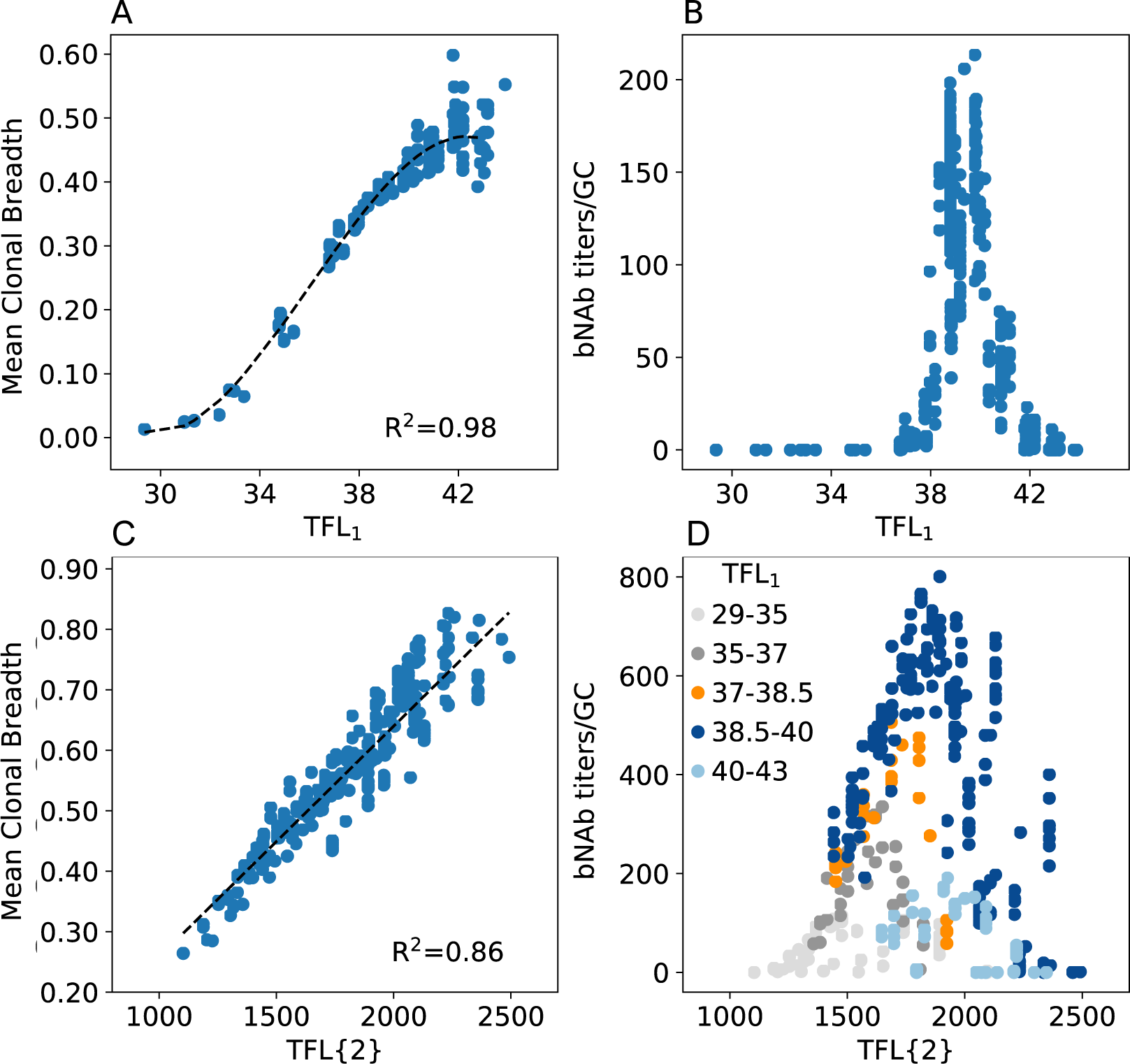
Total frustration level (TFL) of the first single-Ag vaccine immunization (A-B) and two sequential single-Ag immunizations (C-D), versus the mean clonal breadth (A, C) and bnAb titers/GC (B, D). Black dashed lines in (A, C) indicate the polynomial and linear fits used to collapse the data, respectively, resulting in the reported *R*^2^ values. Each dot represents the average output from n=1,000 GCs, and are colored in (D) according to the corresponding value of TFL_1_.

We next wanted to determine if the TFL concept could be used as a metric to predict Ab properties after multiple immunizations, enabling us to determine the optimal way in which to manipulate frustration with time. We hypothesized that the TFL after multiple immunizations is the product of the TFL of each individual immunization. This history-dependent formulation is based on the idea that each new immunization operates on the memory B cells that resulted from the previous immunizations. This more general form of Eqn. 8 is shown in Eqn. 9 for *N* immunizations, where Σ_*j* ≠ *k*_*d*_*jk*_ indicates a summation over the mutational distances between the sequences of all pairs (*N*_*parirs*_) of Ags *j* and *k* administered across all immunizations. Here, *b* is a weight that renormalizes differences in the simulated ranges of the CFL and SFL values; see SI). Terms with a subscript, i, in Eq. 9 refer to each individual immunization (e.g., TFL_1_), and TFL{*N*} refers to the total frustration level after *N* non-GL-targeting immunizations.

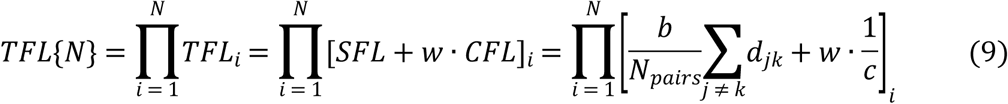

Following the first single-Ag vaccine immunization (Fig. 4A-B), we next simulated a second sequential single-Ag immunization (third immunization overall; Fig. 4C-D). Here, we manipulated frustration by changing concentration (c2) and the average mutational distance (d2) between the current immunizing Ag and both previously-administered Ags (GL-targeting Ag and first vaccine Ag). We find that Eq. 9 is highly predictive of the mean breadth after multiple immunizations (Fig. 4C; *w*_*1*_=*w*_*2*_=24.6, *b*_*1*_=1, and *b*_*2*_=1.6 (see SI)). As before, we also find that the bnAb titers/GC are maximized at intermediate levels of frustration, graphed in Fig. 4D as the net frustration level after all immunizations (i.e., TFL{2}). However, note that different levels of TFL_1_ lead to different peak positions and maximal values of the bnAb titers/GC when the results are graphed against TFL{2}. In the following section, we investigate this behavior, to better understand how best to manipulate frustration over time to maximize bnAb evolution.

### Optimally increasing frustration with successive immunizations maximizes bnAb production via diverse evolutionary paths

To determine the optimal way in which to manipulate frustration with successive immunizations (or time), simulations were performed that varied TFL_2_ at constant values of TFL_1_, namely at a low TFL_1_ (Figs. 5A, D), intermediate TFL_1_ (Figs. 5B, E), and high TFL_1_ (Figs. 5C, F). We find that regardless of the level of frustration imposed upon the immune system in the first immunization, an optimal level of frustration always exists for the second immunization that maximizes the bnAb titers/GC. The origin of the optimality in the bnAb titers/GC after the second immunization is the same as was described earlier for the first immunization. Notably, the optimal TFL_2_ is always higher than the corresponding TFL_1_. This result says that increasing the level of frustration in successive immunizations maximizes bnAb production. This is because after the first immunization, some B cells are likely to have developed moderate to strong interactions with the conserved residues, and a stronger selection force is required for evolving additional mutations that can further focus their interactions on the conserved residues to acquire breadth. A higher level of frustration provides such a selection force to promote bnAb evolution.

**Fig. 5.**
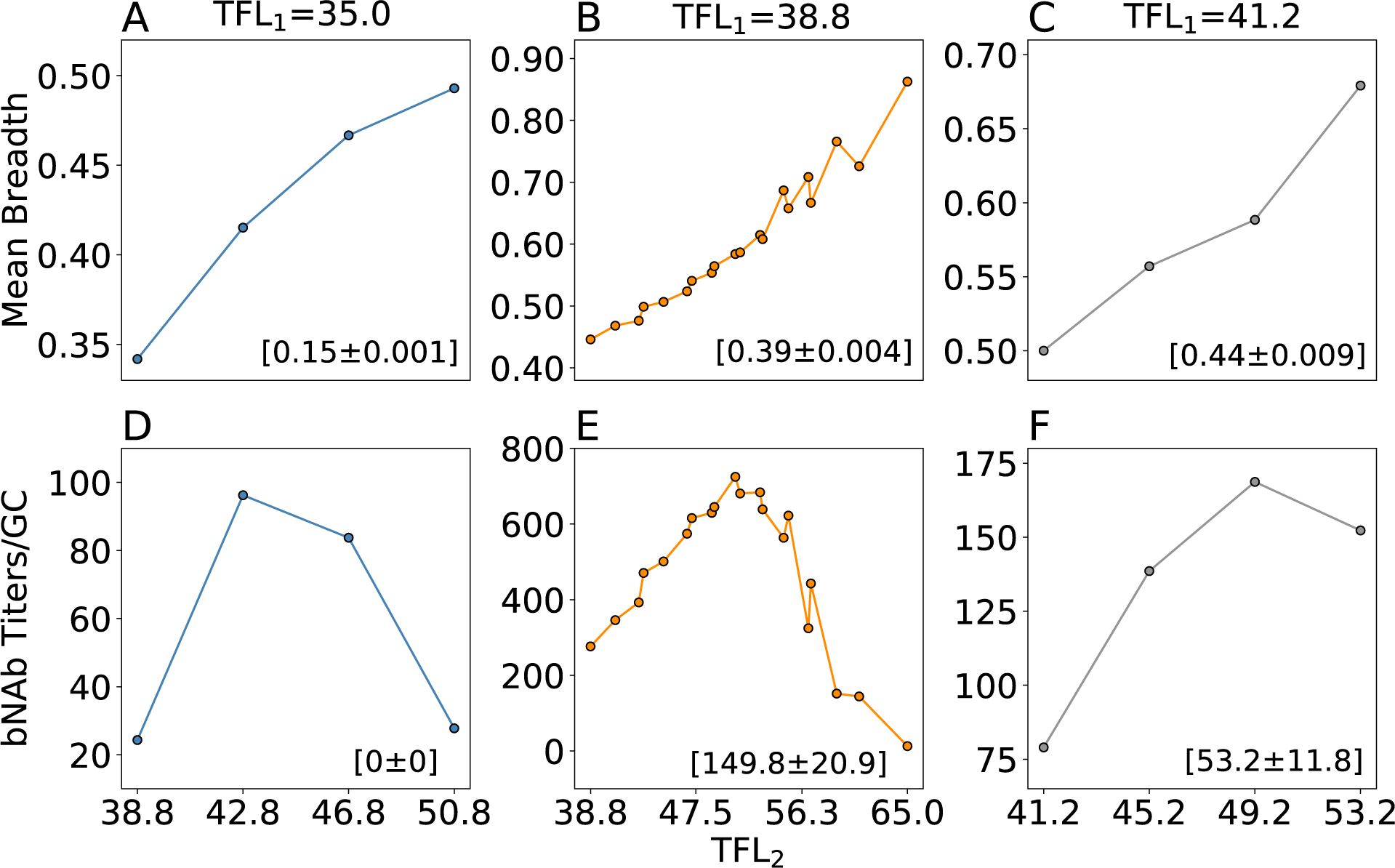
Effect of changing the total frustration level administered in the second immunization (TFL_2_), conditioned on (A) a low total frustration level administered in the first immunization (TFL_1_), (B) an intermediate TFL_1_, and (C) a high TFL_1_, on the mean clonal breadth and bnAb titers/GC after the second vaccine immunization. Values in parentheses indicate the mean clonal breadth and bnAb titers/GC after the first immunization.

Comparing the maximum bnAb titer values in Figs. 5D-F, we find that the highest titers are not only produced at intermediate values of TFL_2_, but at an intermediate value of TFL_1_, which can also be observed in Fig. 4D. Fig. S2 shows the validity of this finding across a wider range of TFL_1_. Taken together, our results imply that each new immunization should be administered at an intermediate level of frustration above the optimum level in the previous immunization to maximize bnAb production. To determine the mechanism underlying such a requirement, we hypothesized that the intermediate value of TFL_1_ resulted in an optimal clonal diversity of memory B cells that provided many potential evolutionary pathways to “success” (i.e., becoming a bnAb; breadth > 0.8) upon the second immunization. To test this hypothesis, we performed a more comprehensive set of simulations that varied TFL_2_ at constant TFL_1_, recording the mutational trajectories of all clones.

Considering only the clonal trajectories that achieved success after the second vaccine immunization, we find that the clonal diversity after the first vaccine immunization was high between a TFL_1_ of 37 and 41 (Fig. 6A). Consistent with our hypothesis above, such an intermediate value of TFL_1_ equal to 39 resulted in the most successful trajectories after the second vaccine immunization (Figs. 5B, D). Similar to our earlier discussion of Fig. 2, this is because at this particular value of TFL_1_, the number of GC cycles – or equivalently frustration – is as high as possible to allow for the most time to make affinity-increasing mutations and thus diversify the clonal population, without incurring major cell death. A lower TFL_1_ resulted in either: 1) low clonal diversity due to too little time to make mutations before the GC reaction ended, providing few pathways for achieving success in the next immunization (TFL_1_<37; Fig. 6C), or 2) high clonal diversity but many of the resulting memory B cells are likely to lead to dead-end evolutionary trajectories that do not evolve bnAbs upon subsequent immunizations at suboptimal levels of frustration (e.g., TFL_1_=37). Values of TFL_1_ higher than the optimum resulted in extensive GC extinction, which restricted the number of successful evolutionary trajectories that could be generated upon the subsequent immunization.

**Fig. 6.**
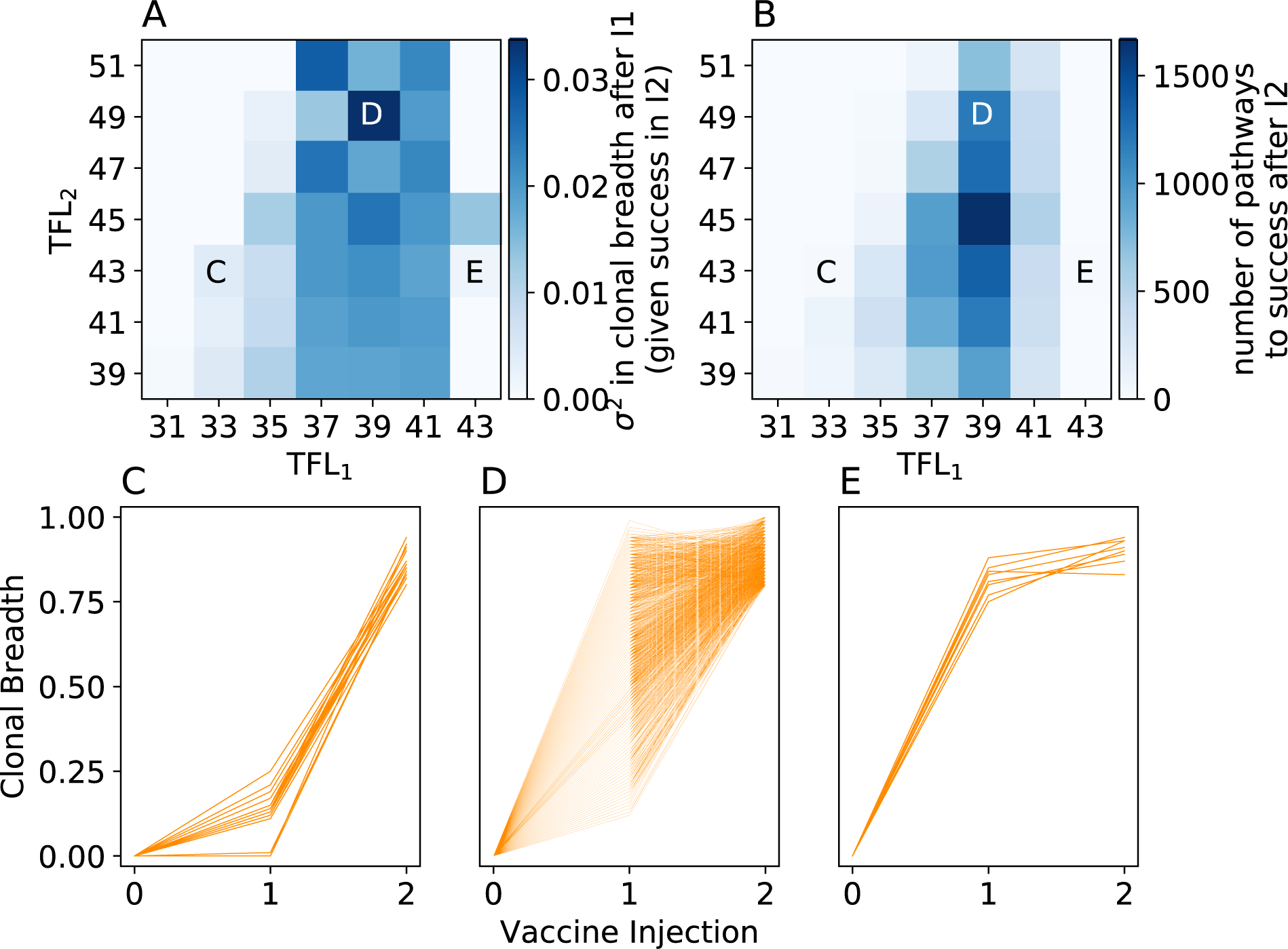
(A) Clonal diversity after the first vaccine immunization (variance in clonal breadth across all n=1,000 GCs), conditioned on ‘success’ in the second immunization (becoming a bnAb; clonal breadth > 0.8). (B) number of successful trajectories after the second immunization. (C, D, E) Mutational trajectories of individual clones across multiple vaccine immunizations and n=1,000 GCs, for (C) a low TFL_1_, (D) an intermediate TFL_1_, and (E) a high TFL_1_. Corresponding points in frustration space are indicated on the top plots.

Finally, we note that in all three scenarios in Fig. 6C-E, upon analyzing the mutational trajectories of clones from one successful GC at a time (here, not conditioned on success), we observe many instances of clonal interference (Fig. S3). We define clonal interference here as instances where the mean breadth of a given clone is below that of another clone after the first immunization, but surpasses the other clone in breadth after the second immunization. This result further emphasizes the importance of ensuring that multiple evolutionary pathways exist for achieving success, as promoted at intermediate frustration levels.

## Discussion

Universal vaccines currently do not exist for diseases like HIV, HCV, influenza, and malaria, in large part because of the high genetic variability of the pathogens that cause these diseases. Broadly-neutralizing antibodies (bnAbs), which target conserved regions of the pathogenic machinery, offer an exciting route to overcome this challenge. While bnAbs have been isolated from a number of patients^54–62^, despite progress, vaccination protocols to elicit them are not available. Recent work identified that varying antigen (Ag) sequence, concentration, or the pattern of administration can modulate bnAb formation by impacting affinity maturation^28–34,63,64^. Efforts to develop systematic strategies to design vaccines against highly mutable pathogens would be greatly aided if a deep mechanistic understanding of the pertinent immunological processes was available. Toward this end, in this study, we employed computational models to investigate the effect of sequential immunization with variant Ags with diverse Ag sequences and concentration on the evolution of bnAbs by AM.

A large number of immunization protocols are made possible by changing both Ag sequence and concentration. Our results show that a simple lower-dimensional representation of this high-dimensional design space is likely predictive of outcome vis-à-vis bnAb production. Specifically, the level of frustration imposed on the immune system upon a single immunization could be formulated as a linear combination of the frustration due to changing Ag sequence and concentration. Then, this level of frustration was multiplied across immunizations to provide the total frustration level (TFL) after multiple immunizations. We found this metric to be highly predictive of the resultant mean antibody breadth and bnAb titers. However, it remains puzzling why such a simple model works for our computational results. While this governing scaling phenomena may not quantitatively translate in an experimental setting, we predict that some combination of the imposed sequence and concentration frustrations will still likely be predictive of the resultant Ab properties. For example, rather than use mutational distance between Ags as a metric of frustration, it may instead be better to employ the difference in their ability to bind different GC B cells as a metric (if such calculations or experiments can be performed accurately).

Our model predicts that an optimal level of frustration imposed on GC reactions upon the first immunization, followed by a temporally increased level of frustration with each subsequent immunization promotes optimal bnAb responses (Fig. 7). An initial optimal level of frustration results in a population of B cells that has the appropriate diversity to subsequently evolve into bnAbs via diverse evolutionary pathways when a stronger selection force to evolve breadth is imposed in subsequent immunizations. Too low or too high a level of initial B cell diversity leads to either extensive GC extinction or dead-end evolutionary paths. Recent advances in controlled Ag release platforms, such as osmotic pumps^63^ or gene delivery vectors^65^, may enable quantitative validation of the role of temporally increasing frustration on bnAb formation. We suggest immunizing with increasingly dissimilar Ags while holding concentration constant, modulating sequence frustration by inserting increasingly diverse amino acids at variable positions adjacent to conserved Ag residues (Fig. 7). Additionally, advances in high-throughput mutagenesis, deep sequencing, and *in vitro* evolution methods (phage, yeast, bacterial, or ribosomal display)^21^ may enable validation of our finding that diverse evolutionary trajectories optimize bNAb responses.

**Fig. 7.**
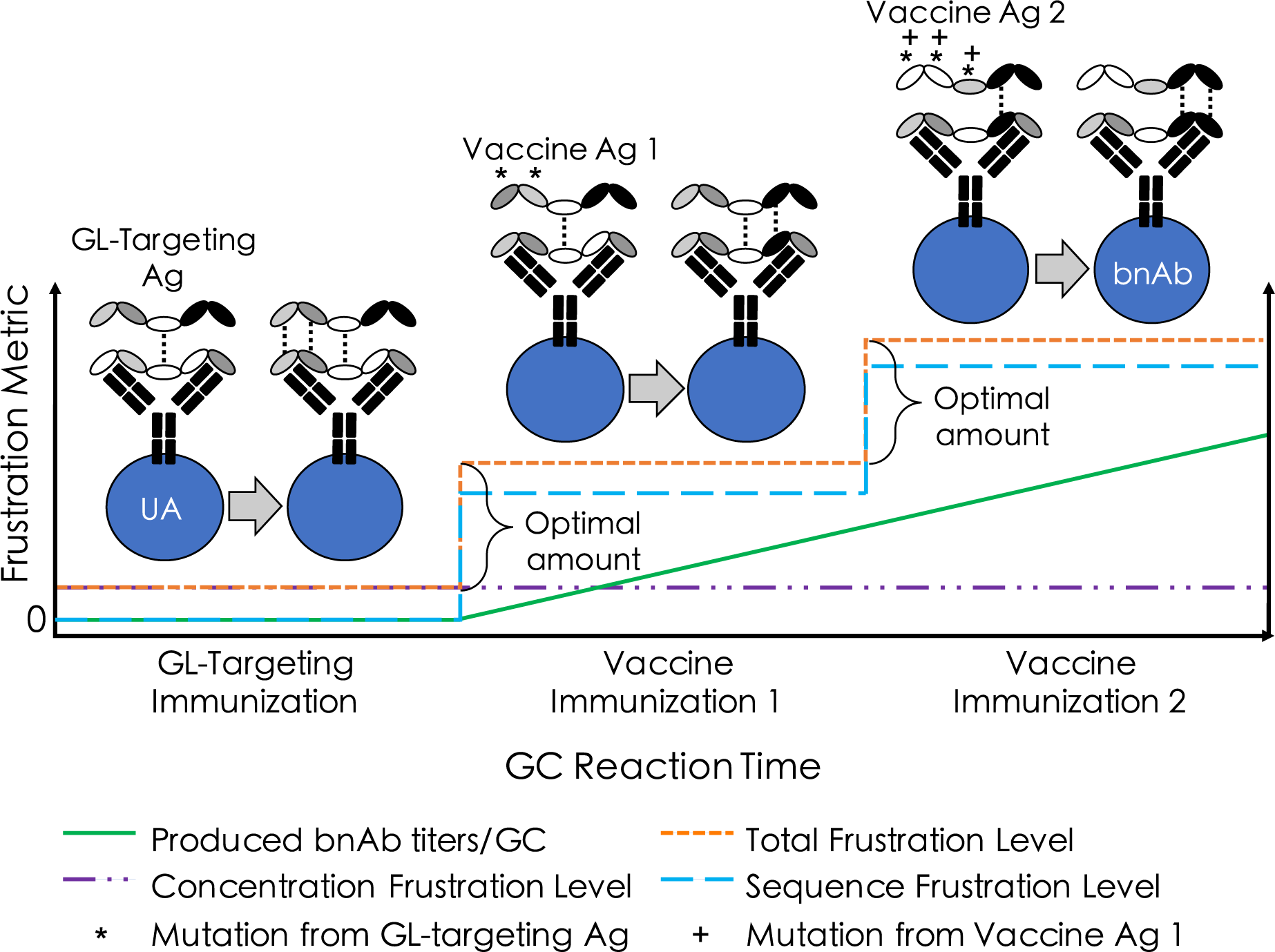
Proposed vaccination scheme. Our model predicts that imposing optimal, temporally increased levels of frustration on GC reactions upon subsequent immunizations (i.e., increasing the ‘total frustration level’; orange line), will optimize bnAb responses (green line). This can be achieved, for example, by immunizing with Ags that are increasingly dissimilar (i.e., increasing the ‘sequence frustration level’; blue line), while holding the concentration of each immunization constant (i.e., keeping the ‘concentration frustration level’ constant; purple line). The total frustration level is the sum of the sequence and concentration frustration levels. A simple schematic of how to increasingly diversify variable Ag residues (white/gray ovals), to guide germline (GL)-targeted unmutated antibodies (UA) towards bnAbs by shifting the focus to conserved Ag residues (black ovals), is demonstrated above the plot.

Like the CD4 binding site (CD4bs) on HIV’s envelope spike protein, the conserved region of the receptor binding site (RBS) on influenza’s hemagglutinin spike protein is smaller in size than the binding footprint of an Ab^66^. Thus, similar to the epitope of CD4bs-directed Abs, the epitope of RBS-directed Abs contains peripheral variable sites in addition to the core conserved sites^66^. Our proposed vaccination approach may therefore be relevant to eliciting bnAbs to the RBS of influenza’s hemagglutinin spike protein. Additionally, in the case of influenza, an intermediate level of frustration in the first immunization may be optimal for another reason, which is accounting for the effects of immunological memory in different individuals. For example, if the vaccine Ag is too different from what a person has been exposed to in the past (i.e., a high level of frustration), then naïve, strain-specific B cells may be more favorably recruited to GCs over cross-reactive memory B cells^67^. After administering the first immunization at an intermediate level of frustration, increasing the frustration in subsequent injections (via differential Ag design, in a similar manner as discussed above for HIV) may transform B cells into true RBS-directed bnAbs^68^.

Our results exhibit an interesting analogy to cognitive learning models^69^, where foundational material is introduced first, followed by more complex material later on. These models, as does our approach, present increasingly difficult material (frustration) over time, allowing many students (B cells) of varying initial skill (breadth) to succeed at each step along the way and eventually pass the test (become a bnAb). This is in contrast to providing complex material first and asking students to take the test right away (akin to a high level of frustration initially), resulting in the success of only a few very bright or lucky students. The idea of increasing frustration with time, particularly sequence frustration, is also similar to how highly mutable pathogens diversify their sequences over time, imposing ever-greater amounts of frustration upon the immune system and creating an evolutionary arms race.

## Acknowledgements

We thank Krishna Shrinivas for helpful discussions. Financial support was provided by the Lawrence Livermore National Laboratory LLC Award #B620960, and by the Ragon Institute of Massachusetts General Hospital, Massachusetts Institute of Technology, and Harvard University.

