## Supplementary material for "Optimizing immunization protocols to elicit broadly neutralizing antibodies": SI

^a^Institute for Medical Engineering and Science, Massachusetts Institute of Technology (MIT), Cambridge, MA 02139; ^b^Harvard-MIT Division of Health Sciences and Technology, MIT, Cambridge, MA 02139; ^c^Department of Chemical Engineering, MIT, Cambridge, MA 02139; ^d^Department of Physics, MIT, Cambridge, MA 02139; ^e^Ragon Institute of MGH, MIT and Harvard, Cambridge, MA 02139; ^g^Department of Chemistry, MIT, Cambridge, MA 02139

*^1^*These authors contributed equally to this work.

_­­­_

***Table S1.*** List of all parameters used in affinity maturation simulations.

| Description | | Parameter | Value |
| --- | --- | --- | --- |
| Somatic hypermutation | Probability of a mutation in CDR or FWR per division | *P_mut_* | 0.14 |
|  | Probability that a CDR mutation is lethal | *P_lethal_* | 0.30 |
|  | Probability that a CDR mutation is silent | *P_silent_* | 0.50 |
|  | Probability that a CDR mutation is affinity-affecting | *P_affinity-affect_* | 0.20 |
|  | Number of total residues | *L* | 46 |
|  | Number of conserved residues | *L_c_* | 18 |
|  | Number of variable residues | *L_v_* | 28 |
|  | Lower boundary for variable residues of seeding B cells | *H_V0, low_* | -0.18 |
|  | Upper boundary for variable residues of seeding B cells | *H_V0, high_* | 0.9 |
|  | Lower boundary for conserved residues of seeding B cells | *H_C0, low_* | 0.3 |
|  | Upper boundary for conserved residues of seeding B cells | *H_C0, high_* | 0.6 |
|  | Lower boundary for residue of B cells | *H_low_* | -1.0 |
|  | Upper boundary for residue of B cells | *H_high_* | 1.5 |
|  | Boundary of energy change due to single-point mutation | *δ* | 1.0 |
|  | Boundary of energy change due to loop insertion mutation | *δ_L_* | 1.0 |
|  | Proportionality factor for loop treatment | *α* | 0.25 |
|  | Mean of shifted lognormal distribution for CDR mutation | *μ* | 1.9 |
|  | Standard deviation of shifted lognormal distribution | *σ* | 0.5 |
|  | Shift/offset of shifted lognormal distribution | *o* | 3.0 |
| Breadth and binding | Pseudo inverse temperature (k_B_T^-1^) | *e_scale_* | 0.08 |
|  | Activation threshold | *E_act_* | 9 |
|  | Breadth binding threshold | *E_th_* | 12 |
|  | Antigen concentration | *c* | varies |
|  | Number of panel Ags to test clonal breadth against | *N_panel Ags_* | 100 |
| GC dynamics | Probability that a B cell is recycled after selection | *P_recycle_* | 0.70 |
|  | Probability that a B cell exits the GC after selection | *P_exit_* | 0.30 |
|  | Fraction of B cells that receive T cell help after binding Ag | *F_help cutoff_* | 0.70 |
|  | Number of B cells that seed a GC | *N_GC founders_* | 10 |

*Choice of* $\alpha$

Immunization with a single Ag has been shown to produce primarily strain-specific Abs; that is, a large fraction of the mutations made by BCRs in response to a single Ag increase binding to variable antigenic sites. However, as our *in silico* vaccination protocols are always preceded by a hypothetical GL-targeting scheme, there should also be a driving force – even for single-Ag administration – towards the conserved residues shared between the vaccine Ag and GL-targeting Ag. We set α to capture both of these facets, namely to achieve approximately equal selection for mutations that increase variable and conserved site binding. For example, given a $\Delta E$ of +1 k_B_T towards a variable site, the overall change in the binding free energy between the Ag and BCR would be a beneficial increase of 0.75 k_B_T ($\Delta E_{overall}=\Delta E+\Delta E_{L}=1-0.25=0.75 k_{B}T)$. Similarly, given a $\Delta E$ of +1 k_B_T directly towards a conserved site, the overall change in the binding free energy would simply be 1.0 k_B_T. The difference of 0.25 k_B_T between these two mutational schemes (summarized in Table S2) is then approximately balanced by the higher probability of mutating in the variable region, due to there being more of these residue types in the simulated epitope (28 variable vs. 18 conserved residues).

***Table S2.*** Example energetic outcomes of different mutation/selection mechanisms against variable vs. conserved antigenic sites, for single-Ag administration.

| Administration type | Mutation Region | $\Delta E$ | $\Delta E_{L}$ | $\Delta E_{overall, BCR-Ag1}$ | $\Delta E_{overall, BCR-Ag2}$ | $\Delta E_{system}$ |
| --- | --- | --- | --- | --- | --- | --- |
|  |  |  |  | $=\Delta E+\Delta E_{L}$ | $=-\Delta E+\Delta E_{L}$ |  |
| Single-Ag | Variable | +1 | - 0.25 | +0.75 | - | +0.75 |
|  | Variable | -1 | +0.25 | -0.75 | - | -0.75 |
|  | Conserved | +1 | - | +1.00 | - | +1.00 |

*Fitting procedures for master curves of mean breadth vs. frustration*

A total of 512 data points, representing individual simulations each of n=1,000 GC trials, were used to construct Fig. 4A (main text). Ten data points were identified as outliers due to a lack of statistics (< 10/1000 GCs succeeded) and removed before fitting took place. The data was fit to a 3^rd^-order polynomial (Equation S1) – for reasons described in the main text – where the TFL1 for each data point was calculated according to Eqn. 8 (main text). We note that while we used a 3^rd^-order polynomial to fit the data in this case, other functional forms may also have been appropriate (e.g., a sigmoidal fit).

$$mean breadth={A\cdot TFL1}^{3}+B\cdot{TFL1}^{2}+C\cdot TFL1+D$$

$$={A\cdot\left[ d1+w_{1}\cdot\frac{1}{c1} \right]}^{3}+{B\cdot\left[ d1+w_{1}\cdot\frac{1}{c1} \right]}^{2}+C\cdot\left[ d1+w_{1}\cdot\frac{1}{c1} \right]+D (S1)$$

The value of $w_{1}$was determined to be that which maximized the *R^2^* correlation value of the data set, resulting in a singular value of 24.6. While this implies that changes in c1 have a larger impact on the TFL than changes in d1 (and thus on downstream Ab production/properties), this finding is highly dependent on the chosen values for *e_scale_* and *E_act_* (Eqn. 4 (main text); Table S1), and on the use of Eqn. 4 itself to characterize the probability of B cell survival via Ag internalization. The final values of the constants A, B, C, and D are listed in Table S3.

For the second sequential immunization, 331 points were used to construct Fig. 4C (main text), where 10 points were again excluded as outliers. Many single-immunization simulations were performed solely to fill out the curve in Fig. 4A (main text), thus there are fewer data points for the second immunization/Fig. 4C (main text). Also, some immunization 1 conditions resulted in complete GC collapse, thus no GCs were initiated upon the second immunization. In this case, we fit the data to a linear form, because for all TFL2 values tested, the minimum energy needed to achieve breadth against the panel Ags was easily exceeded, eliminating the bottom of the sigmoidal-like curve. Maximizing the *R^2^* correlation value of the data set resulted in a singular value for $w_{2}$ of 15.4, a factor of approximately 1.6 below that of the first immunization. The primary reason for this is because the SFL for the second immunization was higher on average than that of the first immunization by close to the same value (~1.75). The higher average SFL in the second immunization was a function of how Ags 1 and 2 were designed relative to one another, specifically to avoid overlapping mutations. Thus, in immunization 2, the impact of the SFL on mean breadth – relative to the CFL – increased compared to in immunization 1. To take such sampling effects into account and be able to compare the TFL across different immunizations (i.e., keep the weights the same; w_1_=w_2_=24.6), we added a correction factor in front of the SFL term in Eqn. 9 in the main text (*b*; for immunization 2, *b*=1.6). As both the CFL and SFL terms were essentially multiplied by this value, this did not change the relationship between the two sources of frustration or the *R^2^* correlation value of the data set.

***Table S3.*** Information on the fitting procedures used to collapse the data in Figs. 4A and 4C (main text).

| Immunization | Number of data points for fitting | Number of data points excluded as outliers | Type of fitting | *w* in Eqns. 8, 9 (main text) | *b* in Eqns. 8, 9 (main text) | Fitting Constants (Poly: A, B, C, D;  Linear: A, B) |
| --- | --- | --- | --- | --- | --- | --- |
| 1 | 512 | 10 | 3^rd^-order polynomial | 24.6 | 1.0 | -0.00040, 0.048, -1.66, 18.85 |
| 2 | 331 | 10 | Linear | 24.6 | 1.6 | 0.00040, -0.12 |

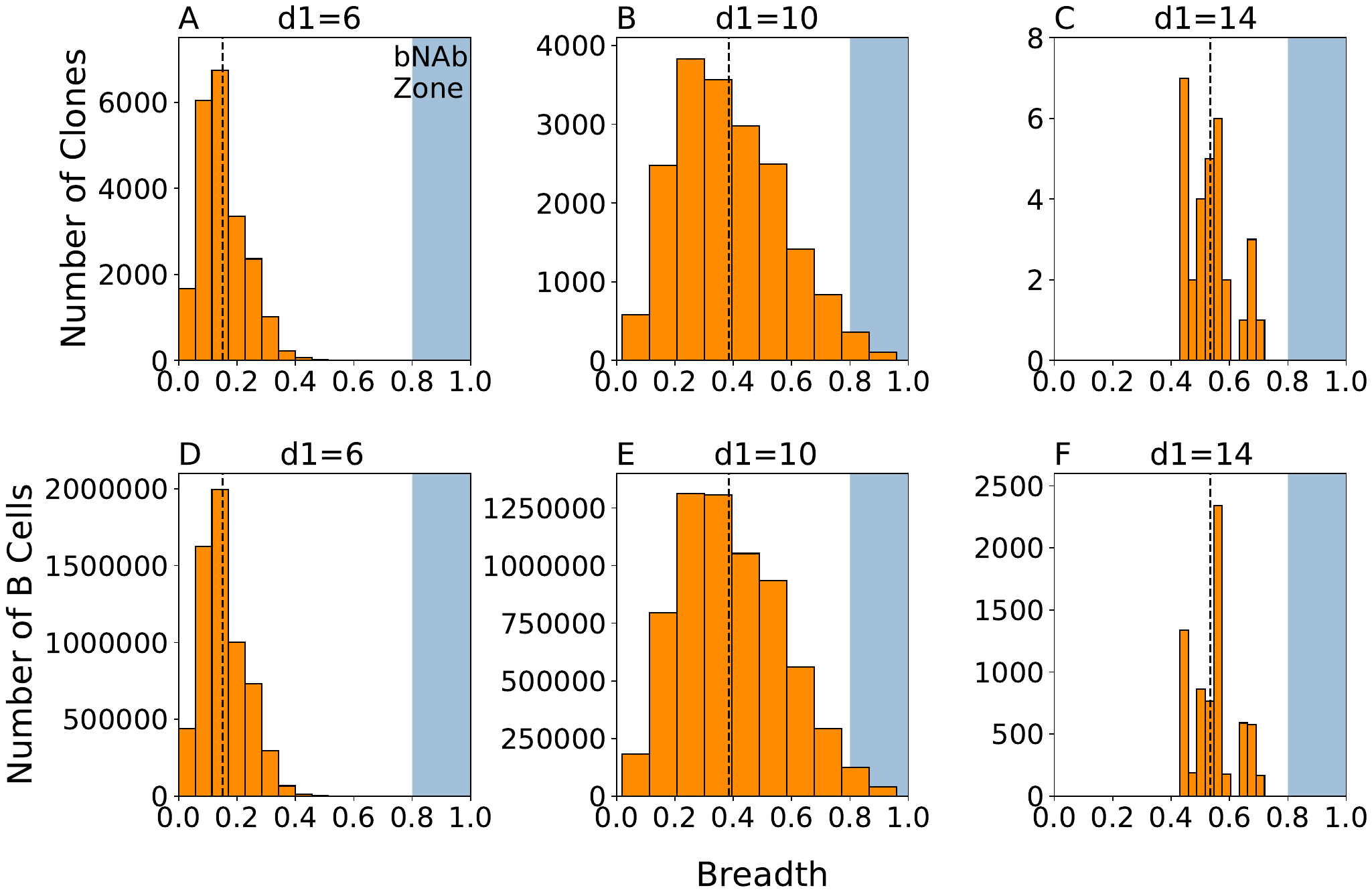

***Figure S1.*** Breadth distributions for (top) all clones and (bottom) all B cells in all clones produced for three different vaccination settings – low frustration (A, D), medium frustration (B, E), and high frustration (C, F). Shaded blue regions and black dashed lines are as described in Fig. 3 (main text). Due to the large number of GCs we analyzed (1,000) for each vaccination setting, the resultant mean breadth and bNAb titers/GC are the same whether we calculate it across all clones or, considering the size of each clone, across all B cells within all clones.

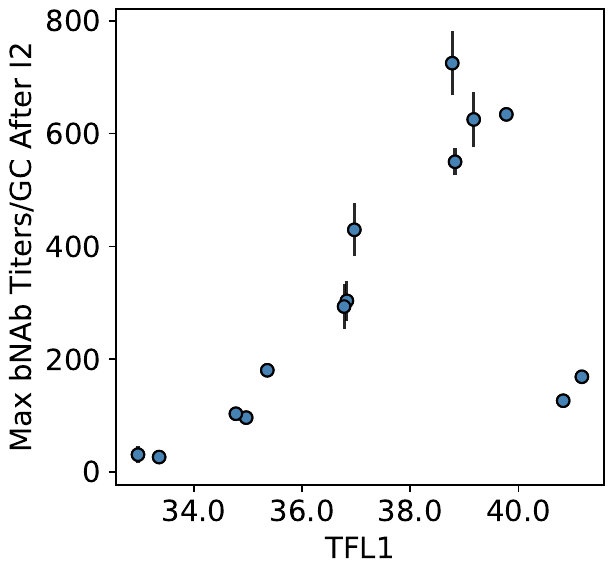

***Figure S2.*** Maximum bNAb titers/GC that can be achieved after immunization 2, for different values of the total frustration level in immunization 1 (TFL_1_).

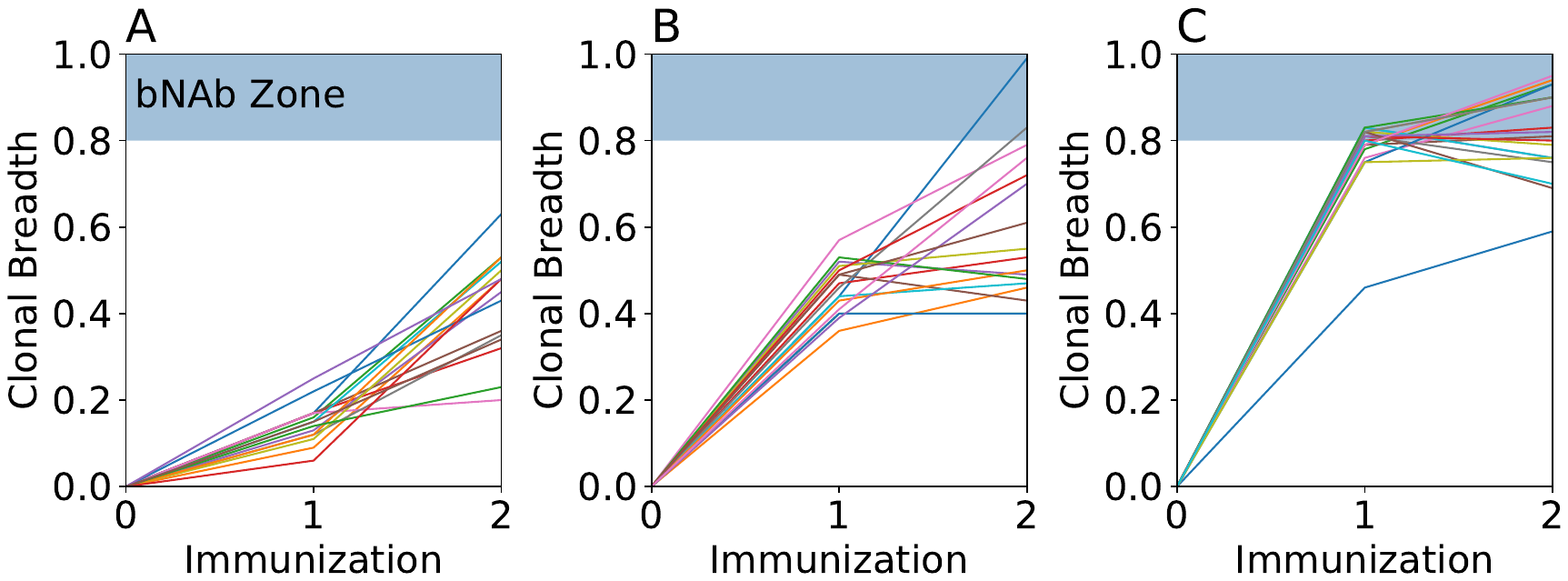

***Figure S3.*** Clonal interference. Mutational trajectories of individual clones from a single successful GC (i.e., simulation trial) after multiple vaccine immunizations. Immunization conditions correspond to those in Fig. 6 (main text): (A) TFL_1_=33, TFL_2_=43; (B) TFL_1_=39, TFL_2_=49; (C) TFL_1_=43, TFL_2_=43. Here, the requirement has been relaxed that the clones must achieve ‘success’ (i.e., a clonal breadth above 0.8 after two vaccine immunizations; see main text).
